# Studies on the Biosorption Potential of Copper by *Rhizopus arrhizus* Biomass

**DOI:** 10.1101/2020.07.13.201566

**Authors:** Jyoti Chauhan, Ishan Saini, Prashant Kaushik

## Abstract

Due to rapid industrialization and global urbanization, heavy metal pollution of water reservoirs is a severe environmental threat. Moreover, the removal of these heavy metal ions of copper from the wastewaters using conventional methods are costly, time taking, and less effective. Whereas, biosorption proved as a better alternative technique over the traditional methods for the removal of metal ions from the water bodies. Biological materials or biosorbents have been used for the adsorption of metal ions from the aquatic system. Therefore, Rhizopus *arrhizus* (living biomass) for the biosorption of copper (Cu) metal was used as biosorbent in the present investigation. The pH and temperature at which biosorption occurs are critical. In case of *R. arrhizus* the maximum adsorption was recorded at pH 7.0, and a temperature of 35°C.Whereas, the maximum adsorption capacity (Q value) of 100 mg biomass of *Rhizopus arrhizus* was observed as 94.46 % at 80 ppm concentration. The maximum adsorption capacity of 200 mg biomass of test fungi was reported as 97.32 % at the metal concentration of 80 ppm. Maximum Q value (biosorption capacity, i.e. mg metal per g biosorbent) was 37.785 mg/g at an 80-ppm concentration in case of 100 mg biomass. In the case of 200 mg biomass, the maximum Q value was 19.464 mg/g observed at 80 ppm concentration. Overall, the present study showed that 100 mg and 200 mg *Rhizopus arrhizus* biomass acts as an excellent adsorbing material for the adsorption of Cu metal ions.

## 1. Introduction

Presence of heavy metal pollutants in the water bodies has become a severe environmental problem [1]. Organic pollutants and heavy metals are regularly reported for crossing their threshold levels in the ecosystem this happens because of the industrial discharge and other human activities that later lead to buildups [2]. Although, heavy metal like copper (Cu) in lower concentrations can be beneficial, at a higher concentration, these metal ions become injurious and act as ecosystem pollutants [3]. The US Environment Protection Agency (EPA) defined Cu as a micronutrient and also as a toxin. It is harmful to brain and kidneys, and also causes other health issues like low blood pressure, gastrointestinal distress, liver cirrhosis and fetal mortality [4]. Excessive level of copper in drinking water results in stomach cramps and diarrhea.

Remediation of these heavy metals ions requires proper care and great attention. Remediation of heavy metals by physical and chemical means is highly expensive and produces a large amount of toxic waste in comparison to other conventional treatment methods [5]. Bioremediation has been proven environment-friendly and less expensive and beneficial for the elimination of metal ions from contaminated water bodies through the microbial uptake [6]. The biosorption (sorption of metallic ions from solutions by live or dried biomass) offers an alternative to the remediation of industrial effluents. Biological materials have the potential to accumulate heavy metal ions from wastewater through metabolically mediated or physico-chemical pathway of metal ion uptake. Therefore, this activity mainly depends on the accumulation of metal ions inside the living cells or adsorbing the metal ions by components of the cell wall [7].

Microorganisms such as bacteria, algae, yeast, and fungi are largely utilized as biosorbent for the biosorption of heavy metal ions from the aquatic system as these can cope the adverse conditions bind with the metal cations due to the presence of negative charge and anionic structures on their cell surfaces. Biosorbents are prepared from waste biomass or naturally abundant biomass [8]. Low cost, high efficiency, minimization of chemical or biological sludge, no additional nutrient requirement, biosorbent regeneration,the possibility of recovery of metal ions, carried out at the contaminated site, and environment friendly are the impressive properties that make the biosorption process quite advantageous as compared to conventional treatment methods [9].

Biosorption process has some disadvantages such as biosorbent gets completely saturated when all the interactive metal sites are occupied, and the improvement of biological process becomes limited as the cells are not metabolizing [10]. Fungal species are served as a potent and eco-friendly tool for the process of biosorption because of high yield of biomass, and they can be chemically and genetically modified [11]. Fungal species are less sensitive to the variation in pH, temperature, and aeration [12].

Therefore, fungal biomass is often chosen as biosorbents for the removal of heavy metal ions [13]. In this direction, it was found that the red alga (*Ceramium virgatum*) is effective and alternative biomass for the removal of Cd(II)ions from aqueous solution [14]. Anayurt et al. [15] used fungal (*Lactarius scrobiculatus*) biomass for the biosorption of Pb(II) and Cd(II) from aqueous solution and reported that recovery of metal ions from this fungal biomass was higher than 95% using 1M HCl and 1M HNO_3_. Umrania [16] reported that microbial biomass could be used for the decontamination of heavy metal ions from wastewaters. Keskinkan et al. [17] used a submerged hydrophytic plant *Myriophyllum spicatum* for the biosorption of Cu and zinc (Zn).

Transportation of metal ions across the cell membrane mainly depends on the metabolic activity of the cell. Only viable cells are capable of this type of biosorption. Cell walls of microbial biomass, mainly composed of polysaccharides, proteins and lipids have abundant metal-binding groups such as carboxyl, sulphate, phosphate and amino groups [18]. Biosorption by living organisms comprises of two steps: First, independent metabolism binding of metals to the cell walls and second, metabolism dependent intracellular uptake, whereby metal ions usually are carried along the cell wall [3]. Holan and Volesky [19] reported that *Rhizopus* sp. is capable for the removal of heavy metals from a different substrate. *Rhizopus arrhizus* has been used for the removal of metal ion from the environment either by complex formation of metal ions with organic acids produced by the fungi or by the adsorption of metal ions to the component of the fungal cell wall. Examples from the literature related to the fungal biosorbents and their uptake efficiency is provided in the Table 1.

**Table 1.** Fungal biosorbents and their uptake efficiency.

| Metal ion | Biomass/biosorbent type | Metal uptake (mg/g) | Reference |
| --- | --- | --- | --- |
| Cu (II) | *Aspergillus niger* | 9.53 | Dursun et al. [20] |
| Cu (II) | *Aspergillus terreus* | 180 | Gulati et al. [21] |
| Fe | *Rhizopus delamar* | 15.7 | Tsekova and Petrov [22] |
| Pb | *Aspergillus versicolor* | 30.6 | Cabuk et al. [23] |
| Cr (VI) | *Lentinus sajor* | 18.9 | Arıca and Bayramoglu [24] |
| As (III), Hg (II), Cd (II), Pb (II) | *Penicillium canescens* | 26.4, 54.8, 102.7, 213.2 | Say et al. [25] |
| Ni | *Penicillium chrysogenum* | 82.5 | Haijia et al. [26] |
| As (III), Hg (II), Cd (II), Pb (II) | *Pencillium purpurogenum* | 35.6, 70.4, 110.4, 252.8 | Say et al. [27] |
| Cd (II), Zn (II), Pb (II) | *Pencillium simplicium* | 52.50, 65.60, 76.90 | Fan et al. [28] |
| Cr (VI) | *Pleurotus ostreatus* | 20.71 | Arbanah et al. [29] ) |
| Cu (II), Ni (II), Zn (II), Cr (VI) | *Pleurotus ostreatus* | 8.06, 20.4, 3.22, 10.75 | Javaid et al. [30] |
| Ni (II), Pb (II) | *Phanerochaete chrysosporium* | 55.9, 53.6 | Ceribasi and Yetis [31] |
| Pb (II), Ni (II), Cr (VI) | *Saccharomyce cerevisiae* | 270.3, 46.3, 32.6 | Ozer and Ozer [32] |

The following two objectives were taken:

(a) The ability to live biomass of *Rhizopus arrhizus* to adsorb Cu metal solutions at different pH and temperature;

(b) The Cu (as cupric sulphate) concentrations at different solutions on the biosorption capacity of living biomass of *Rhizopus arrhizus*.

## 2. Materials and Methods

### 2.1. Preparation of Fungal Biomass, Cu Solution, and Adsorption of Cu by *Rhizopus arrhizus*

Around 1 kg of soil samples were gathered from an organic qualified agriculture field in Meerut, Uttar Pradesh, India. After that, the dilution plate method was applied for the isolation of soil fungi from soil samples as defined previously [33]. For the experimental study of Cu biosorption, *Rhizopus arrhizus* was selected. Potato dextrose agar (PDA) medium was used for the preparation of a pure culture of test fungi. Cu as cupric sulphate was used to investigate the potentiality of *Rhizopus arrhizus* biomass for the metal adsorption. Stock solutions of Cu metal ion stock solutions were prepared for the concentrations of 10 ppm, 20 ppm, 40 ppm and 80 ppm concentration of the metal solution. To examine the capability of Rhizopus arrhizus (living biomass) for the sorption of Cu (cupric sulphate), as explained by Bhole et al. [34] with slight modification was adopted. 50 ml of 10 ppm solution of Cu was taken into each of 6 flasks. Likewise, the same trend was followed for the 20ppm, 40ppm, and 80ppm concentrations of Cu After that, fungal biomass was added as illustrated in Figure 1. Further, kept on a rotatory shaker for 15-20 minutes.

**Figure 1.**
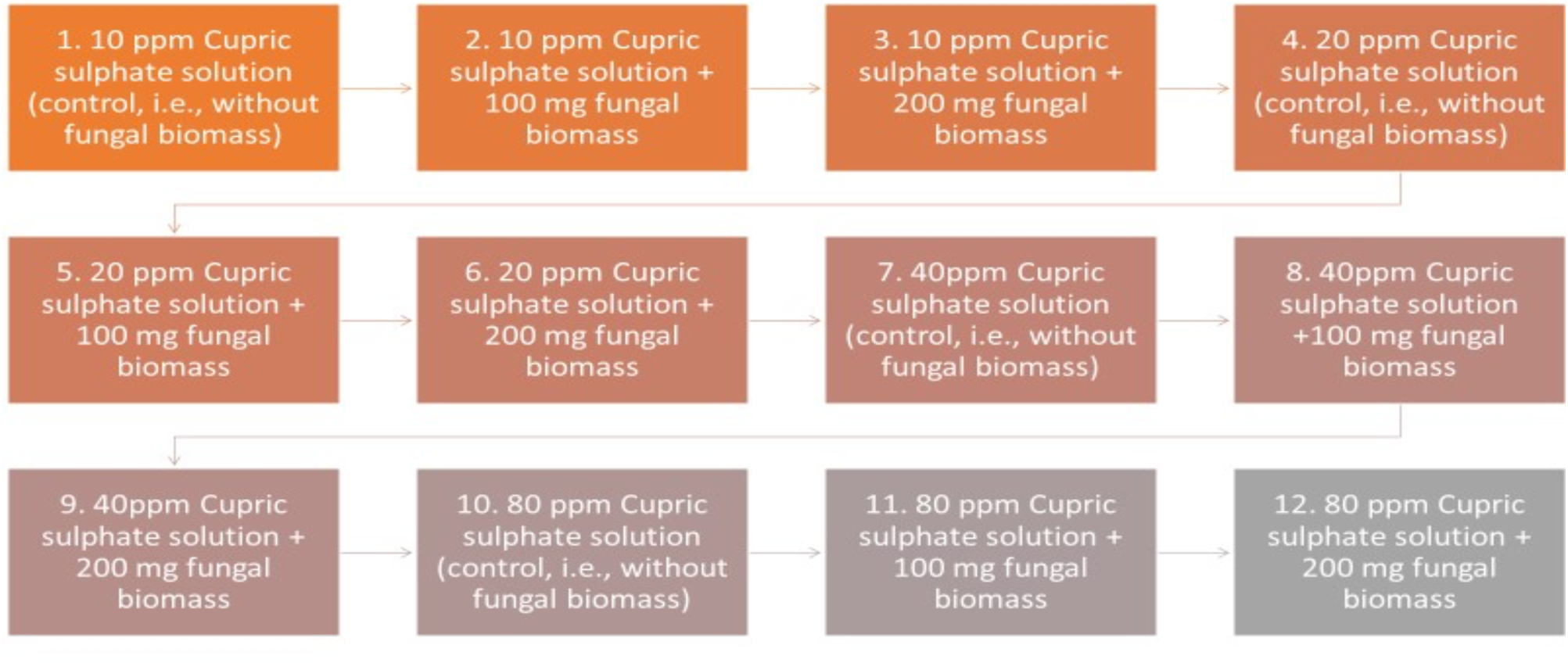
*R. arrhizus* biomass (mg) along with the different concentrations of cupric sulphate (ppm) studied as three replicates in the biosorption study.

After a contact period of 15 minutes, the biomass of fungi was separated by filtering the mixture using Whatman filter paper No. 40. After that, Cu in the supernatant was examined at a wavelength of 640-650nm following the steps mentioned in Figure 2.

**Figure 2.**
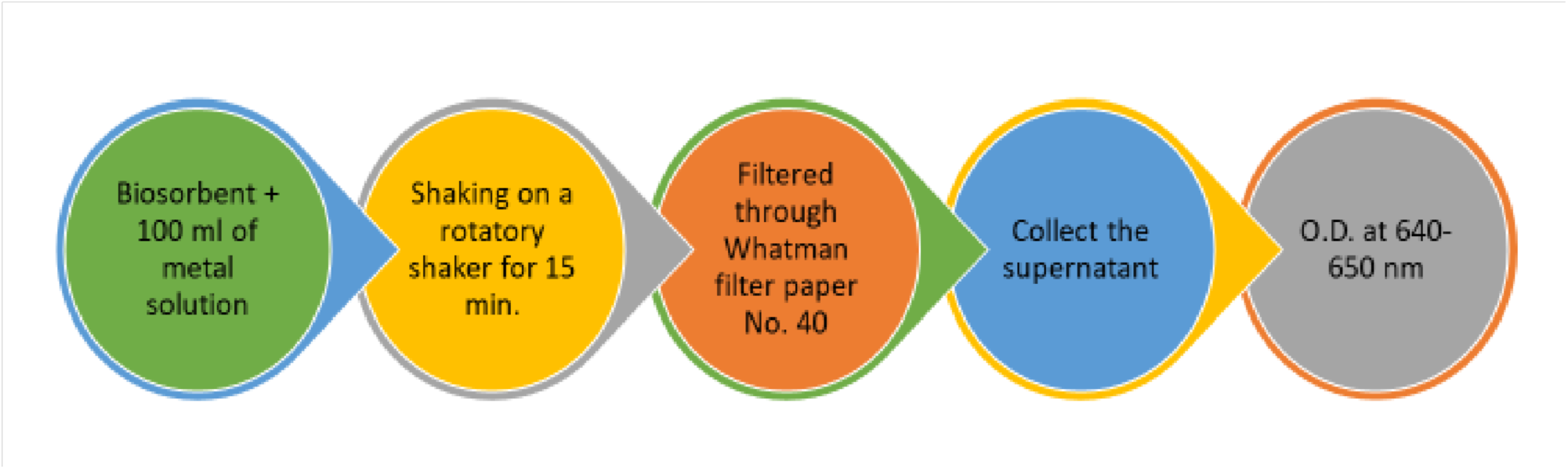
The representation of the steps followed to determine the biosorption potential of *R*.*arrhizus*.

The amount of metal-bound by the biosorbent was calculated based on the formula defined by Hussein *et al*. [35].

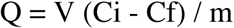

Where,

Q is the metal (Cu Preparation of fungal biomass, Cu solution, and adsorption of Cu by *Rhizopus arrhizus* (living biomass). Where Q is the rate of uptake (mg Cu per g biosorbent), V is the liquid sample volume (ml), Ci is the initial concentration of Cu in solution (mg/l), Cf is the final concentration of Cu in solution (mg/l), and m is the amount of the added biosorbent on a dry basis (mg). Likewise, biosorption efficiency (R %) of the particular biomass can be calculated as:

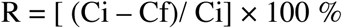

### 2.2. Effect of pH and Temperature

The impact of hydrogen ion concentration on the biosorption of copper metal ion was evaluated by the test solution (that contains 100 mg/l and 200 mg/l initial concentrations) of copper at different levels of pH (3, 5, 7, 9, and 11). Whereas, the temperature (25°C ± 1), the concentration of biomass was kept constant during the study. Temperature effect on the copper adsorption was determined at the 20°C, 25°C, 30°C, 35°C, and 40°C respectively with 100mg/l and 200mg/l initial concentration of copper. During the study, the concentration of biosorbent and pH were maintained constant.

## 3. Results and Discussion

### 3.1. Effect of pH

pH value directly influenced the degree of dissociation of the functional groups present on the cell surface of bioadsorbent and metal ions solubility (Figure 3). The uptake of metal ions increases with pH value (from pH value 3.0 to 7.0) (Figure 4). This is due to the increased attraction sites to metal ions (positively charged), and hence, negatively charged ligands significantly exposed, for the maximum copper biosorption was observed at 7.0. The removal of copper metal ions was about 26.5% at a pH of 3.0 and 78.6% at pH 7.0 at the initial concentration of 100 mg/l. (Figure 3). In the case of the initial concentration of 200 mg/l,the removal efficiencies of copper metal ions were found to be 27.4 % to 82.5 % at pH 3.0 to 7.0.In both cases, the approximately three-time reduction was observed with the decrease in pH from 7.0 to 3.0 (Figure 3).

**Figure 3.**
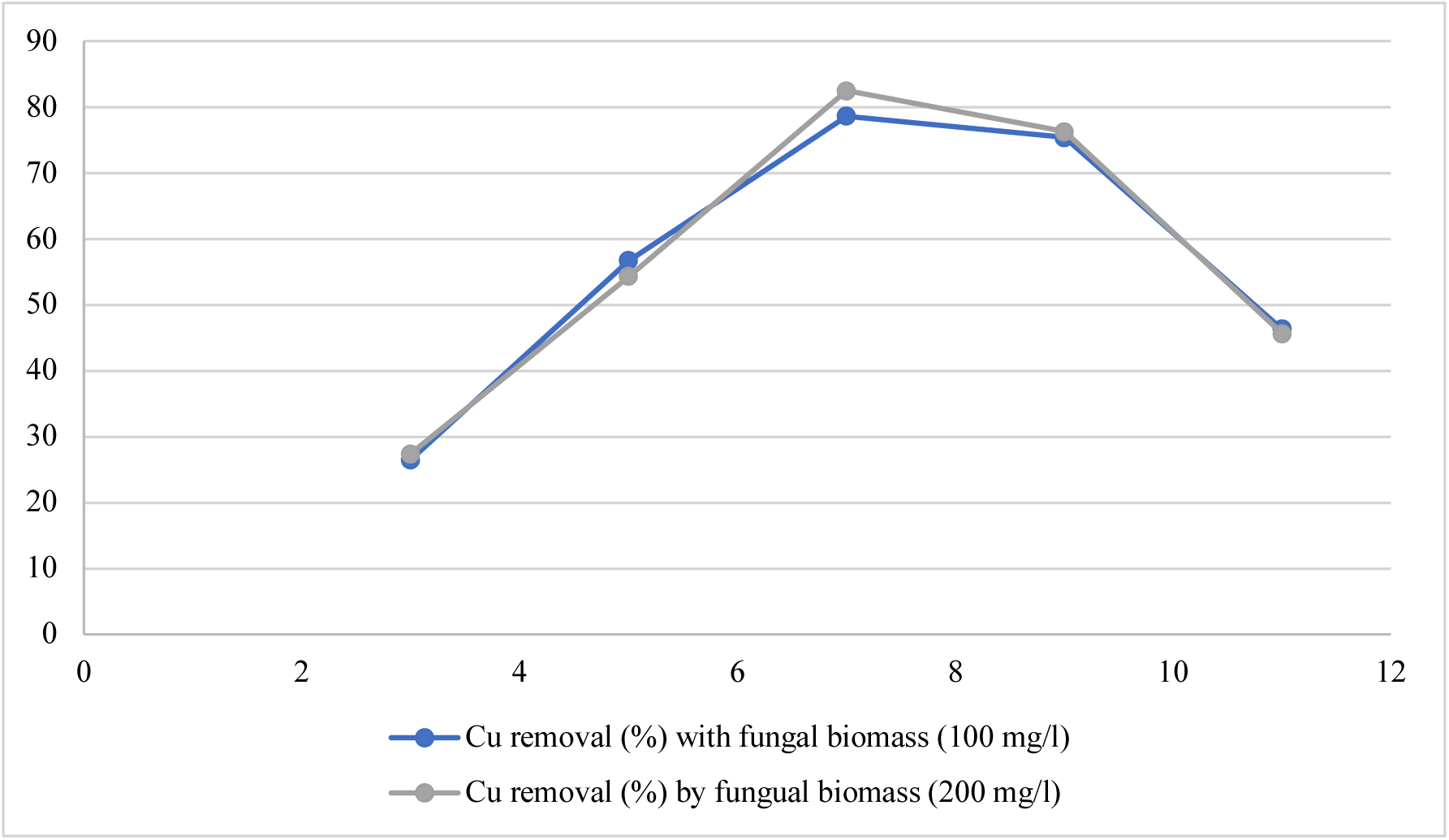
Copper removal efficiency in percentage (y-axis) at different pH (x-axis).

**Figure 4.**
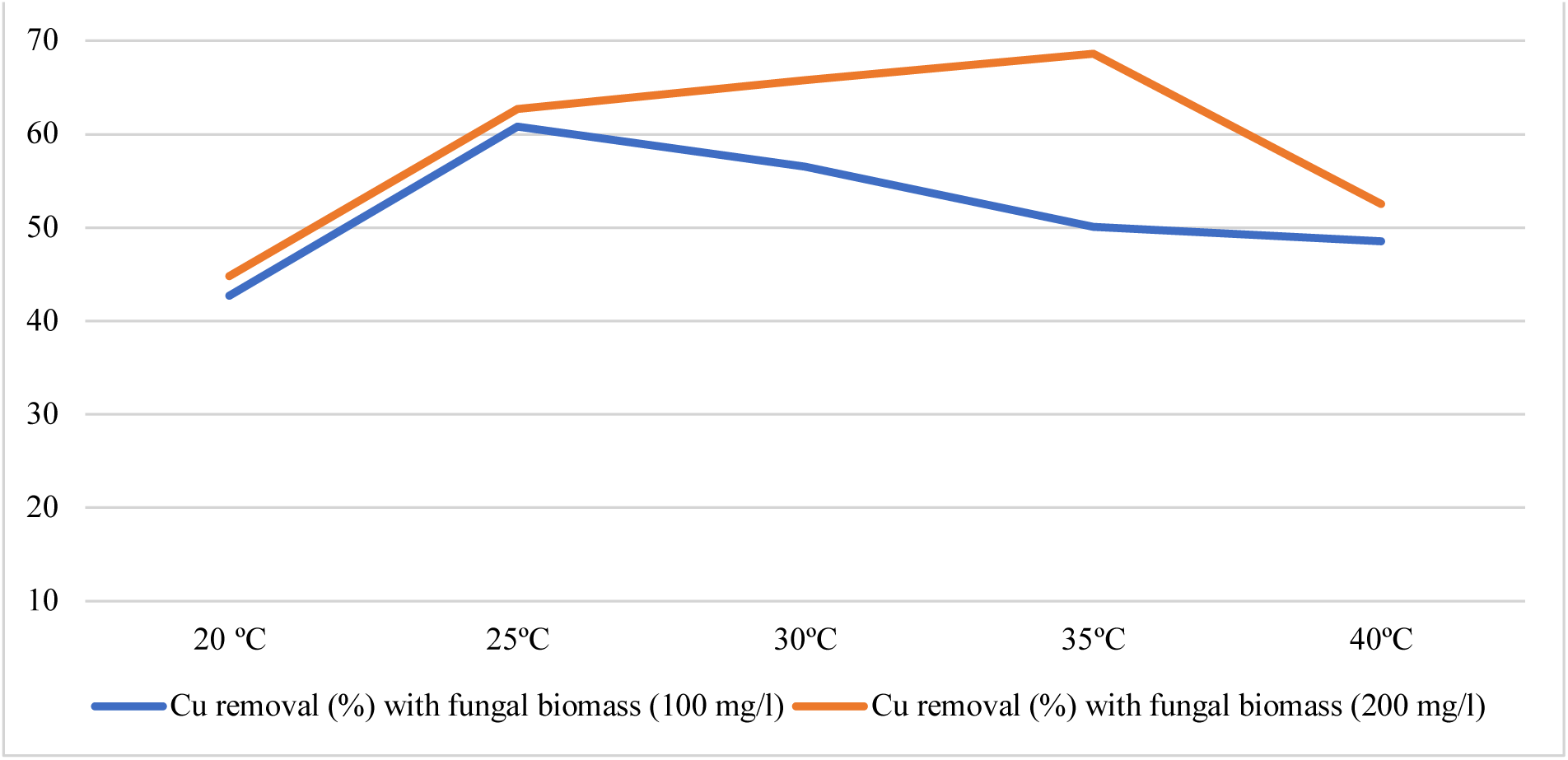
Copper removal efficiency in percentage (y-axis) at different temperature (x-axis).

The net negatively charged groups present on the surface of biosorbent is the primary operating force for metal ion biosorption [36]. Also, the activities of living microorganisms can be hampered at low pH values. With the increased electronegativity of biomass, the force of attraction between the heavy metal cation and biomass increased and due to which the adsorption of heavy metal cation enhanced. The presence of negative charge at the biomass surface promotes the binding of cations or positively charged metal ions. At the low pH of the solution, a competition occurred between the metal ions and proton to bind with the active site present on the surface of biosorbent. Various reports showed that the uptake of metal ions remains low at low pH [37]. It is due to the working of repulsion forces between the metal ions and cell wall of biomass because the cell wall is closely associated with the hydronium ions.

Thus, in the present investigation, copper biosorption by test fungi was observed as a function of pH. But when pH becomes more than 7, the metal hydroxides (Cu(OH)_2_) of metal ions are formed. It was observed that the biosorption efficiency increases from pH 7.0 to 9.0. The increase in pH value indicates two things: (1) the decreased amount of proton, due to this the competition between the metal ions and proton decreases, and (2) greater availability of ligands to binds with the metal ions and as a result biosorption capacity of biomass increased [38]. The biosorption efficiency decreases at the alkaline pH (pH equals 10.0 to 11.0). This is due to the precipitation of metal ions in the form of metal hydroxides. This metal precipitation depends on the concentration of metal ions and pH value but not on the biosorption process. The formation of metal hydroxides or metal precipitation causes interference in the biosorption process as it inactivates the metal ions. Similar results were observed by many workers [39-41].

### 3.2. Effect of Temperature

Biosorption of copper metal ions was studied at a different temperature, i.e., at 20, 25, 30, 35, and 40 °C (Figure 4). Temperature played an essential role during the biosorption of copper (Figure 4). As the temperature of the solution increased, the kinetic energy of copper particles also increased. In case of the initial concentration of 100 mg/l of copper, it was observed that the biosorption of metal ion(copper) progressively increased and maximum biosorption was observed at 25° C temperature than 30 - 40° C (Figure 4). In the case of 200 mg/l initial concentration of copper, it was noticed that maximum adsorption take place at 35°C (Figure 4). Temperature plays a critical role during the biosorption of metal ions as it has a positive and negative effect on the biosorption process [43]. The effectiveness of temperature depends on the system of metal ion biosorption. Gupta et al. [44] reported that the red mud (industrial aluminium waste) worked as a potent biosorbent for the adsorption of Cadmium and Zinc from the aqueous system and also noticed that with the rise in temperature the adsorption decreased. Nuhoglu and Oguz [45] also observed a similar finding for copper biosorption.

### 3.3. Effect of Biosorbent Dose

During this experimentation, it was observed that the removal of copper ions increased with an increasing amount of biosorbent. With the increased biomass, the number of binding sites for metal ion increased. But after some incidences, the increases in biomass the active sites present at the surface of biosorbent become blocked, and as a result of this, the efficiency of biosorbent was observed steady or decreased. The biosorption efficiency of living biomass of *Rhizopus arrhizus* to adsorb Cu from the solutions of different concentrations of Cu solution (as cupric sulphate). Varying amount (100 mg and 200 mg) of living biomass of fungi as biosorbent was used for the biosorption of the Cu from a solution of test metal solutions. The observations were taken after the contact period of 15-20 minutes. Results are presented in Table 2. The living biomass of test fungi is quite useful material for the biosorption of Cu metal ions (Table 2). At the higher concentration (80 ppm) of metal in solution as much as 94.46 % could be adsorbed in 100 mg biomass after the contact period of 15 minutes (Figure 5). While the biosorption of Cu by living biomass of *Rhizopus arrhizus* increased effectively from 27.18 % at 10ppm, 65.95 % at 20 ppm and 86.87 % at 40 ppm, the increase in living biomass of *Rhizopus arrhizus* from 100 mg to 200 mg led to the rise in the biosorption percentage up to 97.32 % (Figure 6). For 100 mg biomass, the Q value was noticed at 80 ppm concentration, i.e., 37.785, followed by 17.374 at 40 ppm, 6.595 at 20ppm and 1.359 at 10 ppm concentrations of metal (Table 2).

**Table 2.** Biosorption of Cu (as cupric sulphate) from the aqueous solution of different concentrations by living biomass (100 mg) of *Rhizopus arrhizus*.

| Amount of Biomass | Initial Conc. of Cu in the solution (ppm) | Final Conc. of Cu in the solution (ppm) | Amount of Cu adsorbed (ppm) | Percentage of Cu adsorbed | Q-value |
| --- | --- | --- | --- | --- | --- |
| 100 mg | 10 | 7.282 | 2.718 | 27.18 | 1.359 |
|  | 20 | 6.810 | 13.19 | 65.95 | 6.595 |
|  | 40 | 5.251 | 34.75 | 86.87 | 17.374 |
|  | 80 | 4.425 | 75.57 | 94.46 | 37.785 |

**Figure 5:**
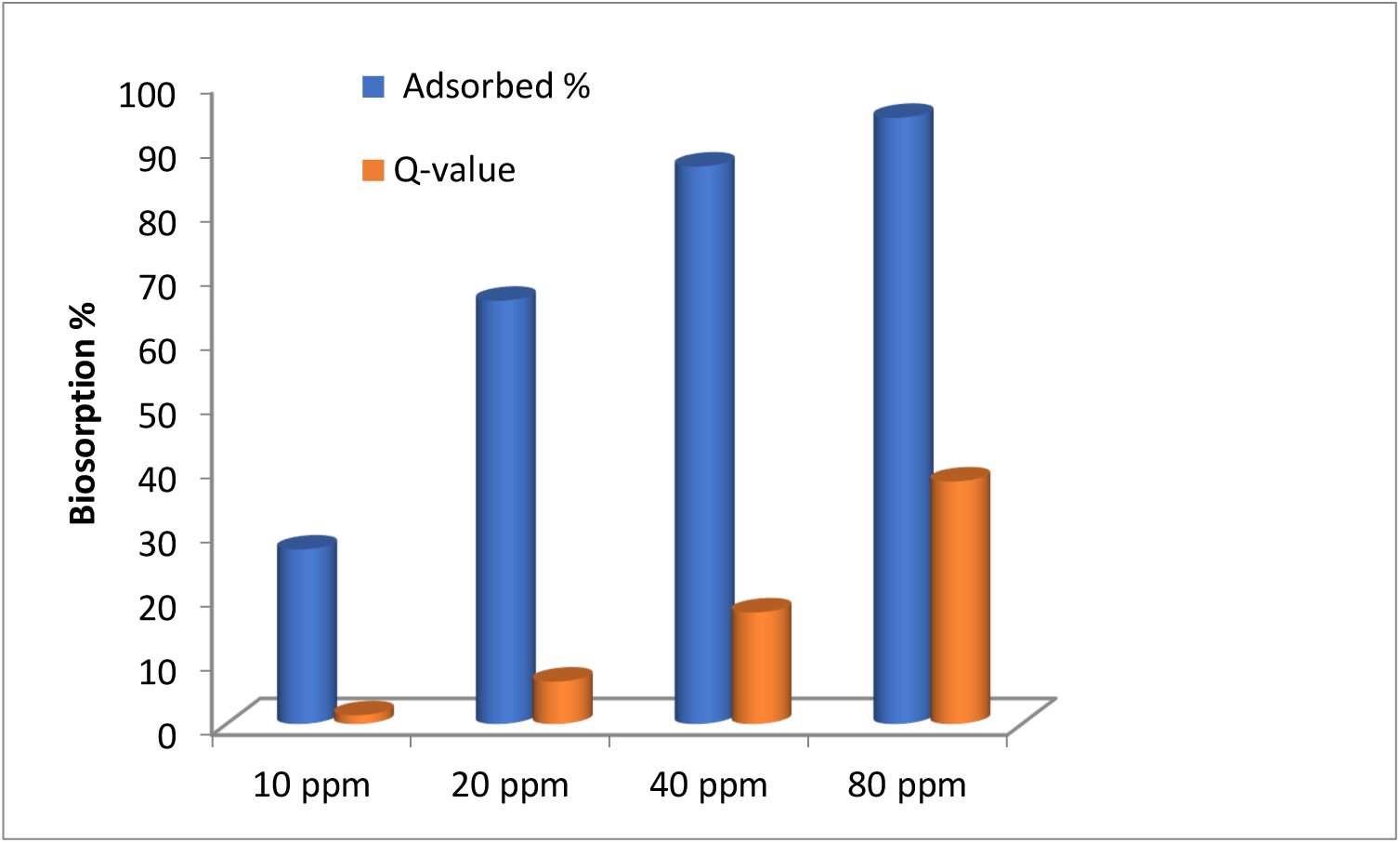
Biosorption profile (%) of Cu by living biomass (100 mg) of *Rhizopus arrhizus*.

**Figure 6.**
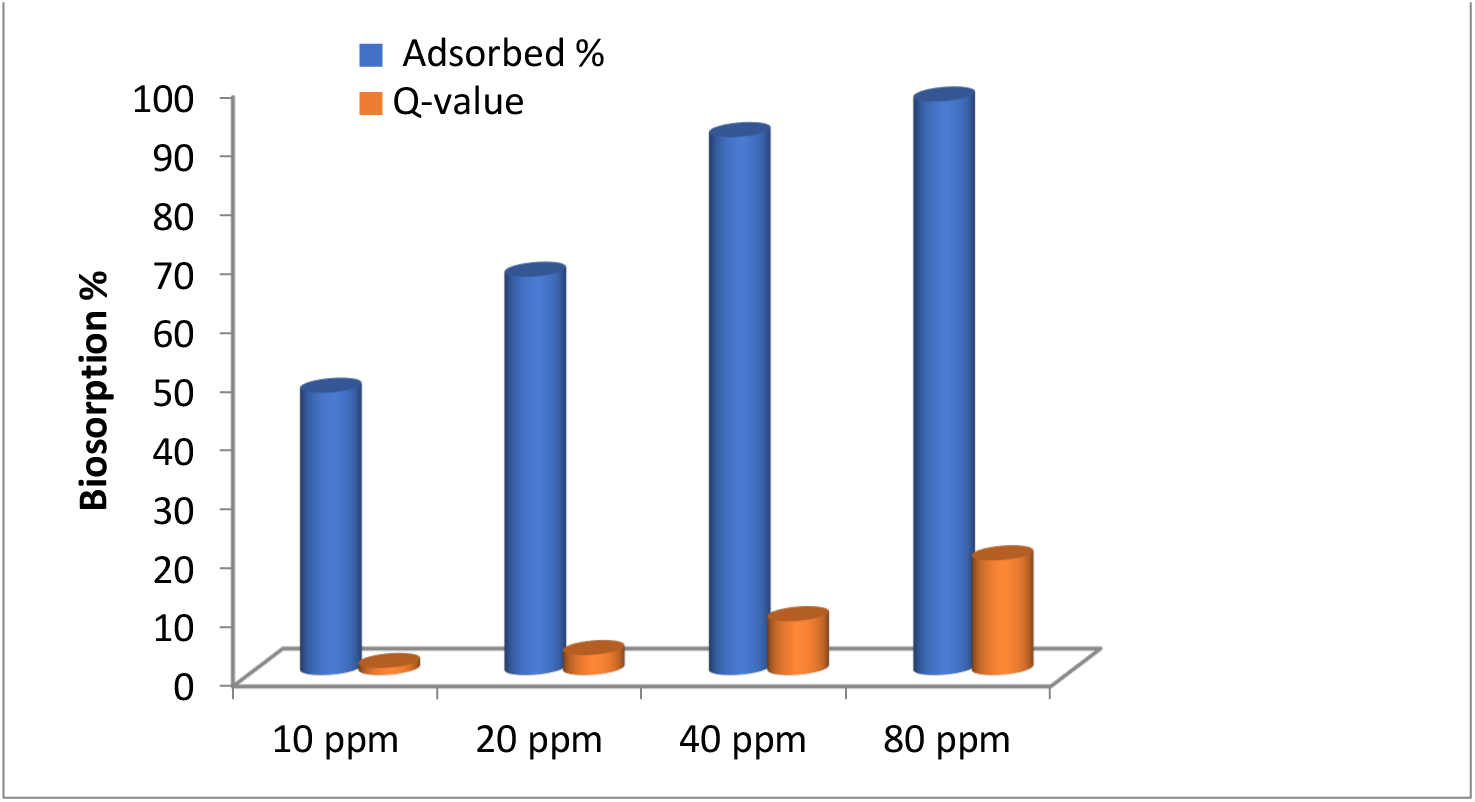
Biosorption profile (%) of Cu by living biomass (200 mg) of *Rhizopus arrhizus*.

About 97.32 % of metal removal was observed from the 80ppm metal concentration by 200 mg fungal biomass (Figure 6). From 10 ppm to 40 ppm, the biosorption of Cu by living biomass of *Rhizopus arrhizus* increased effectively from 47.90 % at 10ppm, 67.59 % at 20ppm and 91.28 % at 40ppm. A glance at Table 3 and Figure 6 showed that in the case of 200 mg biomass, the maximum Q value 19.464 was observed at 80ppm Cu concentration, followed by 9.128 at 40ppm, 3.379 at 20ppm and 1.197 at 10ppm concentrations of a metal solution.

**Table 3.** Biosorption of Cu (as cupric sulphate) from the aqueous solution of different concentrations by living biomass (200 mg) of *Rhizopus arrhizus*.

| Amount of Biomass | Initial Conc. of Cu in the solution (ppm) | Final Conc. of Cu in the solution (ppm) | Amount of Cu adsorbed (ppm) | Percentage of Cu adsorbed | Q-value |
| --- | --- | --- | --- | --- | --- |
| 200 mg | 10 | 5.210 | 4.790 | 47.90 | 1.197 |
|  | 20 | 6.482 | 13.518 | 67.59 | 3.379 |
|  | 40 | 3.487 | 36.513 | 91.28 | 9.128 |
|  | 80 | 2.142 | 77.858 | 97.32 | 19.464 |

The Q values indicated that the boost of fungal had induced a significant decrease in the metallic uptake for every mg of biomass. Previously, it was established that the initial concentrations of metallic ions have a substantial effect on the sorption rate of metals by the fungal biomass [46]. In this direction, it was reported that a saturation point is reached with the increase in biosorbent dose after a specified contact period beyond which desorption occurs, i.e., biosorbent molecules released back into the effluents and this desorption may be followed by resorption [47]. Similarly, Jaikumar and Ramamurthi [48] showed that a rising dose of biosorbent declines the biosorption potential. Chen et al. [49] reported that an increase in specific metal uptake capacity with an increasing initial ions concentration and decreasing in a biosorbent doze. Kahraman et al. [50] reported that with the growing amount of live or dried biomass the biosorption potentiality of biomass firstly increased and after attainment of saturation point, a decrease in biosorption capacity was noticed. The initial increase in the biosorption directly dependent on the dose of biosorbent may be due to the availability of more adsorption binding sites.

## 4. Conclusions

The present study showed that the living biomass of *Rhizopus arrhizus*is an effective biosorbent for the biosorption of Cu metal ions, and it can be used for adsorption-based treatment system of heavy metal ions. Present investigation indicated that the 200 mg (maximum dose of fungal biomass) was the most effective concentration for the Cu biosorption in comparison to the 100 mg. Also, the biosorption efficiency increases with the increasing amount of biomass of test fungi and biosorption of Cu metal ions by living biomass of *Rhizopus arrhizus*is an environmentally friendly and cost-effective technology. The parameter, such as pH of the solution, temperature, and biosorbent dose significantly affects the biosorption process. The concentration of hydrogen ions affects the sorption behaviour of heavy metal ions. pH had a direct effect on the adsorption of copper metal, and maximum sorption of copper metal ions noticed at pH 7.0. The maximum biosorption of copper observed at 35°C. The living biomass of *Rhizopus arrhizus*can be utilized as an effective component for managing heavy metal pollution. Conventional treatment methods for the elimination of metal contaminants from the aquatic system are highly time-consuming and are not eco-friendly. Biosorption is quite an effective alternative to the traditional techniques and is highly efficient for the bio-detoxification of heavy metal pollutants from water bodies and for controlling the environmental pollutants.

## References

1. P.B. Tchounwou, C.G. Yedjou, A.K. Patlolla, and D.J. Sutton: “Heavy Metals Toxicity and the Environment.” EXS. vol. 101, pp. 133–164, 2012.

2. R. Singh, N. Gautam, A. Mishra, and R. Gupta: “Heavy metals and living systems: An overview.” Indian J Pharmacol. vol. 43, no. 3, pp. 246–253, 2011.

3. A.S. Ayangbenro and O.O. Babalola: “A New Strategy for Heavy Metal Polluted Environments: A Review of Microbial Biosorbents.” Int J Environ Res Public Health. vol. 14, no. 1, 2017.

4. F. Pizarro, M. Olivares, R. Uauy, P. Contreras, A. Rebelo, and V. Gidi: “Acute gastrointestinal effects of graded levels of copper in drinking water.” Environmental Health Perspectives. vol. 107, no. 2, pp. 117–121, 1999.

5. V. Masindi and K.L. Muedi: “Environmental Contamination by Heavy Metals.” Heavy Metals. 2018.

6. O.B. Ojuederie and O.O. Babalola: “Microbial and Plant-Assisted Bioremediation of Heavy Metal Polluted Environments: A Review.” International Journal of Environmental Research and Public Health. vol. 14, no. 12, pp. 1504, 2017.

7. I. Timková, J. Sedláková-Kaduková, and P. Pristaš: “Biosorption and Bioaccumulation Abilities of Actinomycetes/Streptomycetes Isolated from Metal Contaminated Sites.” Separations. vol. 5, no. 4, pp. 54, 2018.

8. N.E.-A. El-Naggar, R.A. Hamouda, I.E. Mousa, M.S. Abdel-Hamid, and N.H. Rabei: “Biosorption optimization, characterization, immobilization and application of Gelidiumamansii biomass for complete Pb2+ removal from aqueous solutions.” Sci Rep. vol. 8, 2018.

9. O. Abdi, and M.A. Kazemi: “review study of biosorption of heavy metals and comparison between different biosorbents”. J. Mater. Environ. Sci. vol. 6 no.5: 1386–1399, 2015

10. V. Sameera, D. CH. Naga, B.G. Srinu and T. Y Ravi : “Role of Biosorption in Environmental Cleanup”. J. Microbial Biochem. Technol. R 1, pp. 001, 2011.

11. C. Mulligan, R. Yong, and B. Gibbs : “Remediation alternative treatment option for heavy metal bearing wastewaters: A review”. Bioresource Technology. vol. 53, pp.195–206, 2001.

12. A. L. Leitao : “Potential of *Penicillium* species in the bioremediation field”. Int. J. Environ. Res. and Public Health. vol. 6, pp. 1393–1417, 2009.

13. E.-S.M. El-Morsy: “Cunninghamellaechinulata a new biosorbent of metal ions from polluted water in Egypt.” Mycologia. vol. 96, no. 6, pp. 1183–1189, 2004.

14. A. Sari and M. Tuzen: “Biosorption of cadmium(II) from aqueous solution by red algae (*Ceramium virgatum*): equilibrium, kinetic and thermodynamic studies.” J. Hazard. Mater. vol. 157, no. 2–3, pp. 448–454, 2008.

15. A. Sari, M. Tuzen, D. Citak, and M. Soylak: “Equilibrium, kinetic and thermodynamic studies of adsorption of Pb (II) from aqueous solution onto Turkish kaolinite clay.” Journal of hazardous materials. vol. 149, no. 2, pp. 283–291, 2007.

16. V.V. Umrania: “Bioremediation of toxic heavy metals using acido thermophilic autotrophes.” Bioresour. Technol. vol. 97, no. 10, pp. 1237–1242, 2006.

17. O. Keskinkan, M.Z.L. Goksu, M. Basibuyuk, and C.F. Forster: “Heavy metal adsorption properties of a submerged aquatic plant (Ceratophyllumdemersum).” Bioresour. Technol. vol. 92, no. 2, pp. 197–200, 2004.

18. N. Kuyucak and B. Volesky: “Biosorbents for recovery of metals from industrial solutions.” Biotechnol Lett. vol. 10, no. 2, pp. 137–142, 1988.

19. B. Volesky and Z.R. Holan: “Biosorption of heavy metals.” Biotechnol. Prog. vol. 11, no. 3, pp. 235–250, 1995.

20. A. Dursun, G. Uslu, O. Tepe, Y. Cuci, and H. Ekiz: “A comparative investigation on the bioaccumulation of heavy metal ions by growing *Rhizopus arrhizus* and *Aspergillus niger”*. Biochemical Engineering Journal. vol. 15, pp. 87–92, 2003.

21. R. Gulati, R. Saxena, and R. Gupta : “Fermentation waste of *Aspergillus terreus*: A potential copper biosorbent”. World J. Microbio. Biotechno. vol. 18, pp. 397–401, 2002.

22. K. Tsekova, and G. Petrov: “Removal of heavy metals from aqueous solution using Rhizopus harmful mycelia in free and polyurethane-bound form”. Z. Naturforsch. vol. 57, pp. 629–633, 2002.

23. A. Cabuk, S. Ilhan, C. Filik, and F. Caliskan: “Pb^2+^biosorption by pretreated fungal biomass”. Turk. J. Biol. vol. 29, pp. 23–28, 2005.

24. M.Y. Arica, and G. Bayramoglu: “Cr (VI) biosorption from aqueous solutions using free and immobilized biomass of *Lentinus sajor*-caju: Preparation and kinetic characterization”. Colloids and surfaces A : Physicochemical and Engineering Aspects. vol. 253, pp. 203–211, 2005.

25. R. Say, N. Yilmaz, and A. Denizli: “Removal of heavy metal ions using the fungus *Penicillium canescens”*. Adsorption Sci. Tech. vol. 21, pp. 643–650, 2003.

26. S. Haijia, Z. Ying, L. Jia, and T. Tianwei: “Biosorption of Ni2+ by the surface molecular imprinting adsorbent”. Process Biochemistry. vol. 41, pp.1422-1426, 2006.

27. R. Say, N. Yilmaz, and A. Denizli: “Biosorption of cadmium, lead, mercury, and arsenic ions by the fungus *Penicillium purpurogenum*”. Separation Sci. Tech. vol. 38, pp. 2039–2053, 2003.

28. T. Fan, Y. Liu, B. Feng, G. Zeng, C. Yang, M. Zhou, H. Zhou, Z. Tan, and X. Wang: “Biosorption of cadmium (II), zinc (II) and lead (II) by *Penicillium simplicissimum*: Isotherms, kinetics and thermodynamics”. Journal of Hazardous Materials. vol. 160, pp. 655–661, 2008.

29. M. Arbanah, N.M. Miradatul, and K. Halim: “Utilization of *Pleurotus ostreatus* in the removal of Cr (VI) from chemical laboratory waste”. Int. J. Eng. Sci. vol. 2, pp. 29–39, 2013.

30. A. Javaid, R. Bajwa, U. Shafique, and J. Anwar: “Removal of heavy metals by adsorption on *Pleurotus ostreatus”*. Biomass and Bioenergy. vol. 35, pp. 1675–1682, 2011.

31. I. H. Ceribasi, and U. Yetis : “Biosorption of Ni (II) and Pb (II) by *Phanerochaete chrysosporium* from a binary metal system–kinetics”. Water SA. vol. 27, pp. 15–20, 2001.

32. A. Ozer, and D. Ozer: “Comparative study of the biosorption of Pb (II), Ni (II) and Cr (VI) ions onto *S. cerevisiae*: determination of biosorption heats”. Journal of Hazardous Materials. vol.100, pp. 219–229, 2003.

33. M. Sharma, P. Kaushik, and P. Chaturvedi: “Enumeration, Antagonism and Enzymatic Activities of Microorganisms Isolated from Railway Station Soil.” bioRxiv. pp. 454595, 2018.

34. B.D. Bhole, B. Ganguly, A. Madhuram, D. Deshpande, and J. Joshi: “Biosorption of methyl violet, basic fuchsin and their mixture using dead fungal biomass.” Current Science. vol. 86, no. 12, pp. 1641–1645, 2004.

35. H. Hussein, S. Farag Ibrahim, K. Kandeel, and H. Moawad: “Biosorption of heavy metals from waste water using Pseudomonas sp.” Electron. J. Biotechnol. vol. 7, no. 1, pp. 0–0, 2004.

36. F. Bux, and H.C. Kasan, HC “Comparison of selected methods for relative assessment of sulfate charge on waste sludge biomass”. Water SA. vol 20, pp. 73–76, 1994.

37. P. Kaushik and D.K. Saini: “Silicon as a Vegetable Crops Modulator—A Review.” Plants. vol. 8, no. 6, pp. 148, 2019.

38. P. Kaewsarn, 2002: “Biosorption of copper(II) from aqueous solutions by pre-treated biomass of marine algae Padina sp”. Chemosphere 4, 1081–1085.

39. P. Kalac, M. Ninanska, D. Bevilaque, and I. Staskova, 1996: Concentration of mercury, copper, cadmium and lead in fluting bodies of edible mushroom in the vicinity of mercury smelter and copper smelter. Science Total Environment, 177, 25 1e 258.

40. E. Mehmet, and D. Sukru, 2006: “Removal of heavy metal ions using chemically modified adsorbents”. Journal International Environmental Application & Science, Vol. 1 (1-2): 27–40.

41. P. Putra, A. Kamari, M. Yusoff, F, Ishak, A. Mohamed, N. Hashim, and M. Isa, 2014 “Biosorption of Cu(II) Pb(II) & Zn(II) ions from aqueous solution using selected waste materials adsorption and characterization studies”. Journal Encapsulation Adsorption Science 4, 25e35.

43. Y. Khambhaty, K, Mody, S. Basha, and B. Jha, 2009: “Biosorption of Cr(VI) onto marine Aspergillus niger: experimental studies and pseudo-second order kinetics”. World Journal of Microbiology and Biotechnology Vol 25, 1413–1421.

44. V. K. Gupta and S. Sharma: “Removal of Cadmium and Zinc from aqueous solutions using red mud. Environ. Sci. Technol. 36(16): pp. 3612–3617, 2002.

45. Y. Nuhoglu, and E. Oguz, 2003, “Removal of copper(II) from aqueous solutions by biosorption on the cone biomass of *Thujaorientalis*”. Process Biochem. 38, 1627–1631.

46. P.X. Sheng, Y.-P. Ting, J.P. Chen, and L. Hong: “Sorption of lead, copper, cadmium, zinc, and nickel by marine algal biomass: characterization of biosorptive capacity and investigation of mechanisms.” Journal of colloid and interface science. vol. 275, no. 1, pp. 131–141, 2004.

47. K. Narasimhulu and Y.P. Setty: “Studies on Biosorption of Chromium Ions from Wastewater Using Biomass of Aspergillus niger Species.” J BioremedBiodegrad. vol. 03, no. 07, 2012.

48. V. Jaikumar and V. Ramamurthi: “Effect of biosorption parameters kinetics isotherm and thermodynamics for acid green dye biosorption from aqueous solution by brewery waste.” International journal of chemistry. vol. 1, no. 1, pp. 2, 2009.

49. Z. Chen, W. Ma, and M. Han: “Biosorption of nickel and copper onto treated alga (Undaria pinnatifida): application of isotherm and kinetic models.” J Hazard Mater. vol. 155, no. 1–2, pp. 327–333, 2008.

50. S. Kahraman, D. Asma (Hamamci), S. Erdemoglu, and O. Yesilada: “Biosorption of Copper (II) by Live and Dried Biomass of the White Rot Fungi *Phanerochaete chrysosporium* and *Funalia trogii*.” Engineering in Life Sciences. vol. 5, no. 1, pp. 72–77, 2005.

